# CD4 T cell help prevents CD8 T cell exhaustion and promotes control of *Mycobacterium tuberculosis* infection

**DOI:** 10.1101/2021.02.23.432461

**Authors:** Yu-Jung Lu, Palmira Barreira-Silva, Shayla Boyce, Jennifer Powers, Kelly Cavallo, Samuel M. Behar

## Abstract

CD4 T cells are essential for immunity to tuberculosis because they produce cytokines including interferon-γ. Whether CD4 T cells act as “helper” cells to promote optimal CD8 T cell responses during *Mycobacterium tuberculosis* is unknown. Using two independent models, we show that CD4 T cell help enhances CD8 effector functions and prevents CD8 T cell exhaustion. We demonstrate synergy between CD4 and CD8 T cells in promoting the survival of infected mice. Purified helped, but not helpless, CD8 T cells efficiently restrict intracellular bacterial growth *in vitro*. Thus, CD4 T cell help plays an essential role in generating protective CD8 T cell responses against *M. tuberculosis* infection *in vitro* and *in vivo*. We infer vaccines that elicit both CD4 and CD8 T cells are more likely to be successful than vaccines that elicit only CD4 or CD8 T cells.

## Introduction

Tuberculosis (TB) is a serious global health issue. BCG is the only approved TB vaccine and despite achieving clinically significant protection in infants and toddlers, it has variable efficacy against pulmonary disease, which is the form of disease that leads to transmission (Fine, 1995). More effective vaccines are required to stop transmission and reduce global TB burden. To develop more effective vaccines, we need a better understanding of how *Mycobacterium tuberculosis* (Mtb) evades immunity and how the different elements of our immune system interact to contain and eliminate Mtb infection.

T cells are required for containment of primary Mtb infection and most TB vaccines in development elicit T cell responses (Andersen and Scriba, 2019; Müller et al., 1987; North, 1973). While Mtb infection elicits both CD4 and CD8 T cells, the increased risk of TB in HIV-infected persons highlights a crucial role for CD4 T cells (Bruchfeld et al., 2015; Geldmacher et al., 2010; Geldmacher et al., 2008; Kalsdorf et al., 2009). The essential role for CD4 T cells in immunity to Mtb is supported experimentally by the early mortality of mice lacking CD4 T cells (Mogues et al., 2001). While mice lacking CD8 T cells also die prematurely after Mtb infection, they survive longer than mice without CD4 T cells, which has led to the conclusion that CD4 T cells are more important for protection than CD8 T cells (Mogues et al., 2001). The inability of CD8 T cells to transfer significant protection to immunodeficient mice supports this assertion (Barber et al., 2011). However, there are other interpretations of these data. For example, studies on people with AIDS have demonstrated the importance of CD4 T cells in regulating virtually every arm of the immune response. The contribution of CD4 T cells in generating optimal CD8 T cell and B cell responses is called CD4 T cell help. We hypothesize that CD8 T cells require CD4 T cell help to develop into effector CD8 T cells capable of controlling Mtb infection.

CD4 T cell help enables CD8 T cells to respond more efficiently to viruses and cancer (Borst et al., 2018). CD4 T cells promote CD8 T cell expression of effector molecules such as IFNγ, TNF, granzyme A and B, and facilitate greater migration of CD8 T cells by upregulating several chemokine receptors (Ahrends et al., 2017). The development of CX3CR1^+^ CD8 T cells, noted for their cytolytic function, requires CD4 T cell help (Zander et al., 2019). Helped CD8 T cells express transcriptional networks that promote effector and memory cell development (Ahrends et al., 2017). Indeed, CD4 T cell help during CD8 T cell priming promotes memory development and imprints a quick effector recall response upon challenge (Janssen et al., 2003; Northrop et al., 2006; Shedlock and Shen, 2003; Sun and Bevan, 2003). In contrast, helpless CD8 T cells upregulate inhibitory receptors characteristic of exhausted CD8 T cells (Ahrends et al., 2017).

If CD4 T cells mediate both effector and regulatory functions during TB, the question arises whether the lack of CD4 T cell help compromises CD8 T cell responses and contributes to the susceptibility of CD4 T cell deficient mice. This is a difficult hypothesis to test as it requires dissociating the effector and regulatory functions of CD4 T cells. Nevertheless, other studies have addressed how CD4 T cells affect CD8 T cell immunity during TB. CD8 T cell priming and recruitment to the lung occurs in CD4-deficient mice, although the quantity and quality of Mtb-specific CD8 T cells elicited without CD4 T cells hasn’t been assessed (Caruso et al., 1999; Serbina et al., 2001). During TB, CD4 T cell help promotes CD8 T cell IFNγ production and cytolytic activity, but whether helped CD8 T cells mediate greater protection is unknown (Bold and Ernst, 2012; Green et al., 2013; Serbina et al., 2001). To answer these questions, we compared CD8 T cell responses in infected WT or MHCII KO mice and confirmed our results in a novel adoptive transfer model in which purified CD8 T cells from uninfected mice were transferred with or without Ag85b-specific CD4 T cells into TCRα knockout (KO) mice. We found that helped and helpless Mtb-specific CD8 T cell responses differed in effector molecule and coinhibitory receptor expressions. Importantly, we found that helped CD8 T cells mediate better Mtb control than helpless CD8 T cells, leading us to predict that vaccines that elicit both CD4 and CD8 T cell responses are likely to confer the greatest protection.

## Results

### Mtb-specific CD8 T cell responses are generated independently of CD4 T cells

MHCII KO mice were used to study the CD8 T cell response to Mtb in the absence of CD4 T cell help. The “helpless” CD8 T cells (i.e., from MHCII KO mice) were compared to “helped” CD8 T cells from Mtb-infected C57BL/6 mice (hereafter, WT). MHCII KO mice had significantly higher bacterial burden in lungs and spleens (Figure S1A) and succumbed early after infection as previously described (Figure 1A) (Mogues et al., 2001). To determine whether CD4 T cell help is required to generate CD8 T cell responses in the lung, we measured Mtb-specific CD8 T cell responses for two dominant epitopes, TB10.4_4-11_ and 32A_309-318_ (Carpenter et al., 2016; Sutiwisesak et al., 2020). These CD8 T cell responses were elicited by Mtb infection without apparent delay in the absence of CD4 T cells (Figure 1B). The frequencies and absolute numbers of TB10.4_4-11_ specific CD8 T cells were higher in MHCII KO than WT mice early during infection, which may have been driven by the greater bacterial load in MHCII KO mice (Figure 1B, S1B). CD8 T cells differentiate into KLRG1^+^CD127^−^ short-lived effector cells (SLEC) or KLRG1^−^CD127^+^ memory precursor cells (MPEC) upon activation. The frequencies of SLEC and MPEC CD8 T cells in the lung of WT and MHCII KO mice were similar. However, the frequencies of both SLEC and MPEC TB10.4_4-11_ -specific CD8 T cells were greater in WT mice than in MHCII KO mice (Figure 1C, 1D). These results indicate that CD8 T cell responses to Mtb are generated independently of CD4 T cell help. However, CD4 T cells promote the differentiation of CD8 T cells into SLEC and MPEC cells, consistent with results previously shown in a vaccination model (Ahrends et al., 2017).

**Figure 1.**
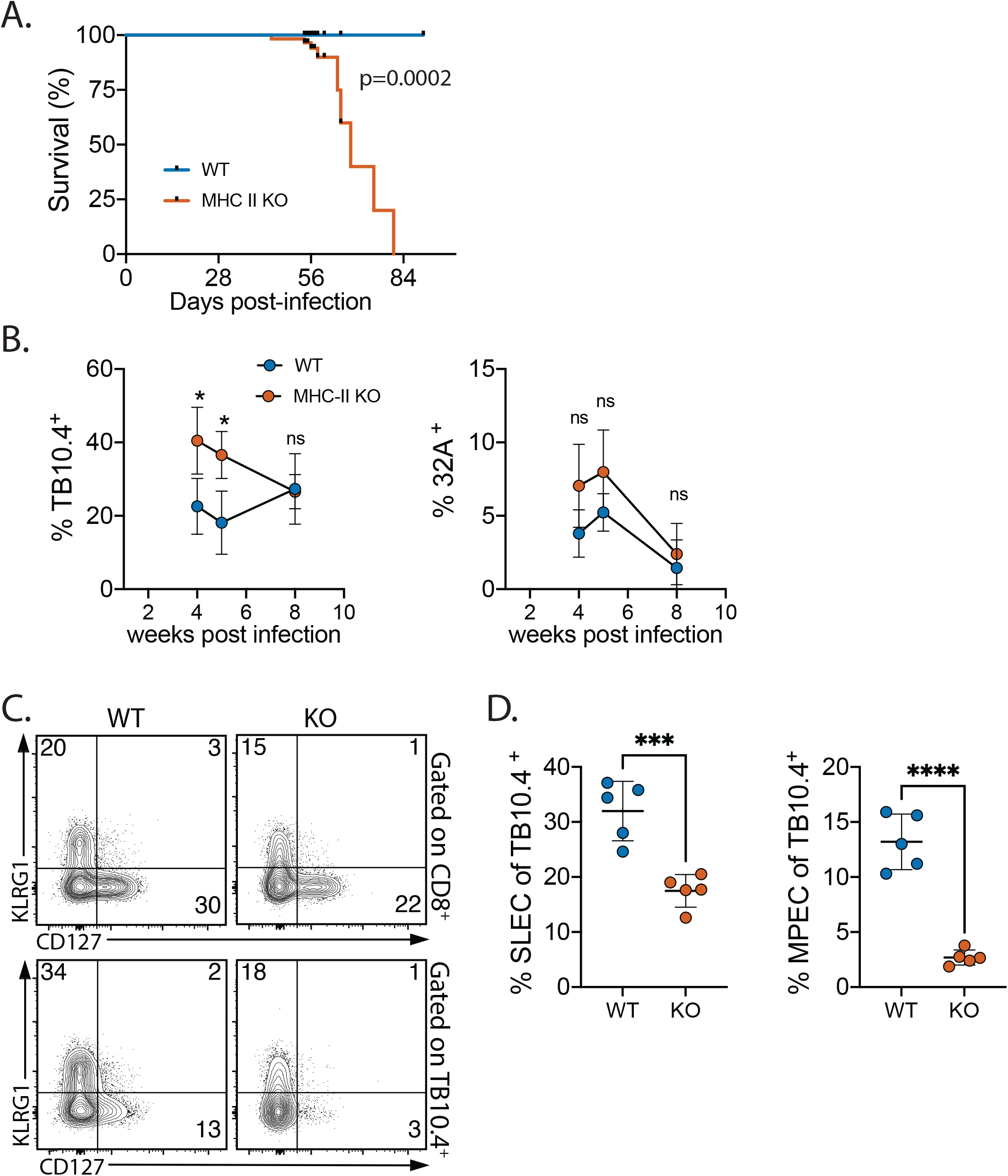
Mtb-specific CD8 T cell responses are generated in MHCII KO mice. (A) Cumulative survivals of WT and MHCII KO mice after infection with aerosolized Mtb. Data were compiled from 9 experiments with 56 WT and 57 MHCII KO mice. The black ticks represent censored subjects (56 WT and 48 MHCII KO mice) that were analyzed at defined time points. The difference between the strains was statistically significant (p=0.0002) as determined by the log-rank test. (B) Lung cells from WT and MHCII KO mice were analyzed at 4, 5, and 8 wpi to determine the frequencies of TB10.4_4-11_ and 32A_309-318_ -specific CD8 T cells. The frequencies of KLRG1^+^CD127^−^ (SLEC) and KLRG1^−^CD127^+^ (MPEC) among total CD8 T cells and TB10.4_4-11_ -specific CD8 T cells was determined 8 wpi by flow cytometry (C, representative data) and analyzed statistically (D). (B, D) Representative data of three independent experiments, 4-5 mice/group. Data represent mean ± SD. Statistical significance was analyzed by multiple t test (B) or unpaired t test (D). p-values: *, p<0.01; ***, p<0.001; ****, p<0.0001. ns, no significant differences.

### Helped vs helpless CD8 T cells: transcriptional signatures of cytotoxic vs. exhausted CD8 T cells

To comprehensively identify differences between “helped” and “helpless” CD8 T cells, we purified lung CD8 T cells from WT or MHCII KO mice 8 weeks post infection (wpi) and performed RNA-seq. We found 1,072 genes differed in their expression between WT and KO CD8 T cells (padj<0.05 and |log_2_ fold-change|*≥*1). Among these genes, 173 were upregulated in WT cells, while 899 genes were upregulated in KO cells (Figure 2A). Among the genes upregulated in WT cells, MHCII molecules were the ones with the largest fold changes. Enrichment analysis using Gene Ontology (GO) terms revealed WT CD8 T cells expressed pathways involved in IFNγ production and cell killing (Figure 2B). These data show that IFNγ production and cytotoxicity by CD8 T cells is augmented by CD4 T cells during Mtb infection. Gene sets associated with NK cell immunity were detected in the enrichment analysis, including expression of NK receptors *Klrk1* (NKG2D) and *CD244* (2B4) (Figure 2B, 2C). These receptors are associated with secretion of multiple cytotoxic granules, indicating enhanced cytotoxicity of helped CD8 T cells (Balin et al., 2018). Since the RNA-seq analysis was performed using total lung CD8 T cells, which contain approximately 50% intravascular cells and could dilute the signal attributable to Mtb-specific CD8 T cells, we changed the criteria to “padj <0.05” without restrictions on fold-change. More genes associated with an effector state were identified as upregulated in WT cells including transcription factors that drive effector CD8 T cell differentiation (e.g., *Tbx21*, *Zeb2*, and *Id2*), activation markers (e.g., *Il2ra*, *CD69*, and *Itgal*), and cytotoxic molecules (e.g., *Gzma* and *Gzmb*) (Figure 2C).

**Figure 2.**
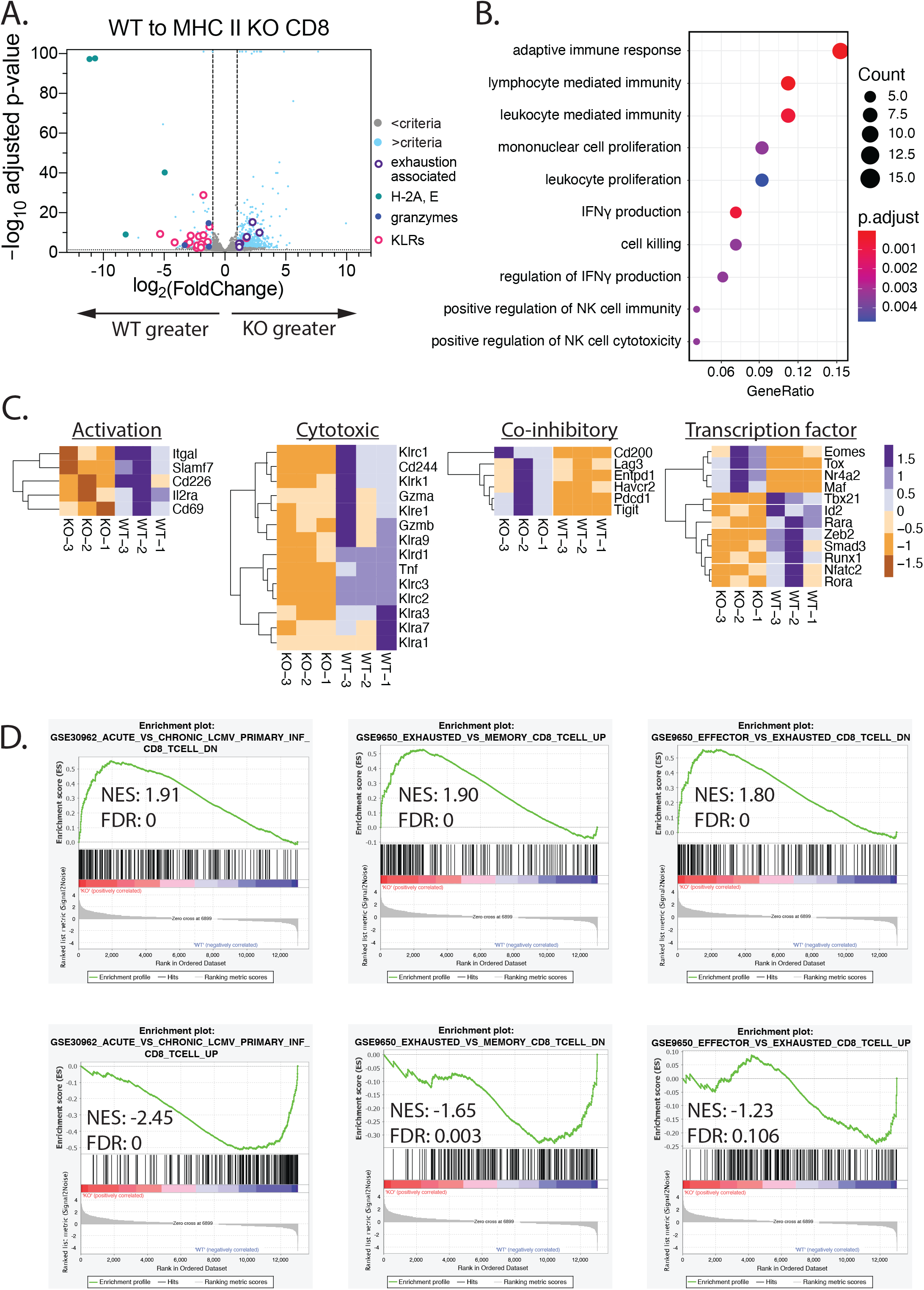
Transcriptional features of WT vs. MHCII KO CD8 T cells. (A) Volcano plot of transcriptome comparison. Transcripts of purified MHCII KO CD8 T cells were compared to WT CD8 T cells from Mtb-infected mice 8 wpi. Blue dots indicate differentially expressed genes (p-adjusted value <0.05 and |log_2_ fold change|*≥*1). (B) Gene Ontology analysis for genes significantly upregulated in WT CD8 T cells. Bubble plot presents top 10 pathways that were most significantly enriched. (C) Heatmap displaying differentially expressed genes in each category. Color indicates row z scores. (D) GSEA. Data were compared to published datasets available at Gene Expression Omnibus (GEO): GSE30962 and GSE9650. NES, normalized enrichment score. FDR, false discovery rate.

Gene set enrichment analysis (GSEA) found that the MHCII KO CD8 T cells upregulated genes associated with T cell exhaustion and were enriched for genes from the signature of CD8 T cells from mice with chronic LCMV infection (Wherry et al., 2007) (Figure 2D, top row). Genes upregulated in KO cells were enriched in genes differentially expressed by exhausted cells compared to effector or memory CD8 T cells (Figure 2D, top row). Indeed, MHCII KO CD8 T cells upregulated many genes encoding inhibitory receptors, including *Pdcd1*(PD-1), *Havcr2* (Tim-3), *Lag3*, and *Entpd1* (CD39) (Figure 2C). Transcription factors *Eomes, c-Maf,* and *Tox* were all upregulated in MHCII KO CD8 T cells (Figure 2C). In contrast, WT CD8 T cells upregulated genes from the effector and memory signatures (Figure 2D, bottom row). Thus, helped WT CD8 T cells express an effector transcriptional program that is remarkable for greater expression of molecules associated with cytotoxicity including TNF, granzyme A and B, and NK receptors. In contrast, helpless MHCII KO CD8 T cells have an exhaustion signature including PD-1, Tim-3, Tox, CD39 and Lag-3.

### Helped vs helpless CD8 T cells: cytotoxic vs. exhausted CD8 T cells

To validate the transcriptional signatures, we performed single cell analysis of CD8 T cells from infected WT or MHCII KO mice using 19-parameter flow cytometry. Since cells located in lung parenchyma are more likely to interact with infected cells, an established approach was used to distinguish intravascular from parenchymal CD8 T cells (Anderson et al., 2014; Anderson et al., 2012). More parenchymal WT CD8 T cells expressed NK receptors including NKG2D and 2B4 than parenchymal MHCII KO CD8 T cells. In contrast, MHCII KO CD8 T cells expressed significantly more co-inhibitory receptors, including PD-1, Tim-3, Lag-3, and CD39, typical of exhausted CD8 T cells (Figure 3A). Differences in transcription factor expression were consistent with our RNA-seq analysis, with a dramatic increase in Eomes expression by MHCII KO CD8 T cells (Figure 3B). WT CD8 T cells expressed higher levels of T-bet and Tcf-1. These differences between WT and MHCII KO CD8 T cells were also identified in total lung CD8 T cells (Figure S2A, S2B).

**Figure 3.**
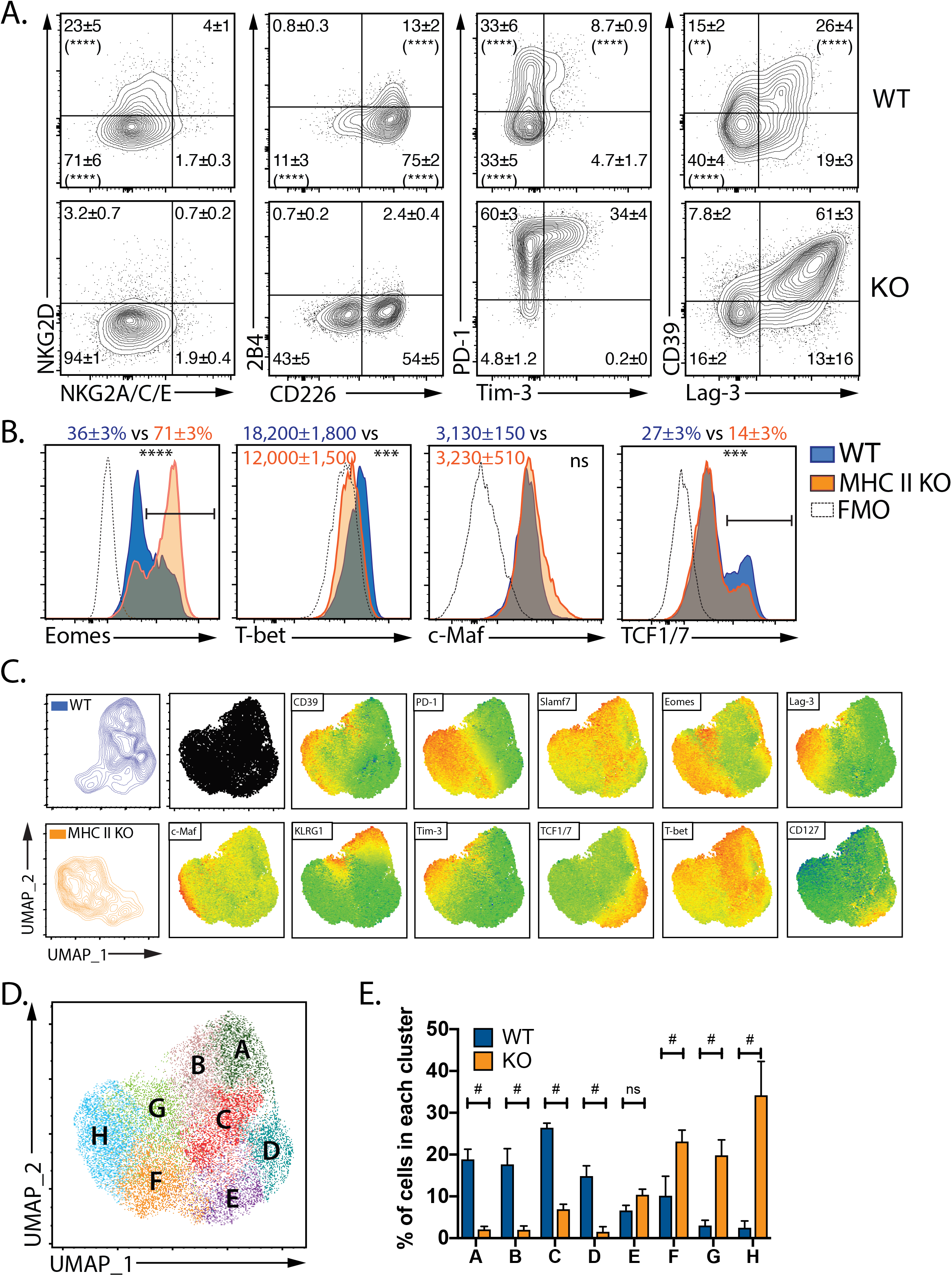
WT CD8 T cells are cytotoxic effectors, while MHCII KO CD8 T cells are exhausted cells. WT and MHCII KO mice were infected with aerosolized Mtb for 8 weeks. Anti-CD45-AF647 was injected intravenously 3 minutes before euthanasia. (A) Lung parenchymal (CD45iv^−^ CD44^+^CD62L^−^) CD8 T cells were analyzed by flow cytometry to determine the frequencies of NK and inhibitory receptors. The mean % ± SD for each quadrant is indicated (n=5/group). The percentage of WT vs. KO cells in each quadrant was compared, and if statistically significant, is indicated in the WT quadrant. (B) The median fluorescence intensity (MFI) or average percentage of cells expressing each transcription factors, ± SD, are indicated. (C) UMAP projections from the flow cytometric analysis of lung parenchymal CD8 T cells were overlayed with expression of the indicated markers. (D) Eight clusters were identified using PhenoGraph and the percentage of each cluster among WT or MHCII KO lung parenchymal CD8 T cells were analyzed statistically. Bars, mean ± SD. (A-E) Representative data of two independent experiments, 5 mice/group. Statistical significance was analyzed by two-way ANOVA (A, E) or unpaired t test (B). p-values: **, p<0.01; ***, p<0.001; ****, p<0.0001; #, p<0.0001. ns, no significant differences.

To facilitate the flow cytometric analysis, UMAP projections were created to visualize differences between the WT and MHCII KO CD8 T cells during Mtb infection (https://arxiv.org/abs/1802.03426). Intravascular and parenchymal CD8 T cells were easily distinguished in the UMAP projection (Figure S2C). In the lung parenchyma, most CD8 T cells were antigen-experienced (CD44^+^), but a minor population of naïve cells (CD62L^hi^CD44^lo^) was also identified. The majority of MHCII KO CD8 T cells localized to the lung parenchyma, perhaps driven by the higher bacterial burden in MHCII KO mice (Figure S2C, S2D).

Antigen-experienced (CD44^+^CD62L^−^) lung parenchymal WT and MHCII KO CD8 T cells were clustered into eight distinct populations on the UMAP projection using PhenoGraph™(Levine et al., 2015) (Figure 3C, 3D). Clusters A-D were dominated by WT CD8 T cells (Figure 3E), which were T-bet^+^Slamf7^+^PD1^lo^, suggesting an effector state (Comte et al., 2017; Kurtulus et al., 2019; Loyal et al., 2020). Cells in cluster A expressed Klrg1, Tcf1, and Eomes. While Tcf1 and Eomes can regulate the generation of memory and exhausted T cells, both populations arise from Klrg1^−^ populations (Angelosanto et al., 2012; Wherry et al., 2007). Thus, Cluster A may represent cells in transition. Cluster B and C expressed T-bet and Slamf7, which defined these as effector cells. Cluster B are terminal effector cells based on their KLRG1 expression. As both KLRG1^+^ and KLRG1^−^ populations expressed NKG2D, NKG2A/C/E, and 2B4, cluster B and C might vary in their differentiation states rather than having distinct effector functions (Figure S2D). Cluster D expressed Tcf1, Eomes, and CD127, consistent with memory precursor cells (see Figure 1, MPECs) (Zhou et al., 2010).

Clusters E-H were predominantly MHCII KO CD8 T cells and were distinguished by their expression of PD-1 (Figure 3C, 3E). The PD-1^low^TCF1^+^Eomes^low^ phenotype of cluster E was typical of exhausted progenitor CD8 T cells, while cluster H appeared to be terminally exhausted CD8 T cells based on expression of the inhibitory receptors Tim-3, Lag-3, and CD39, and the transcription factors Eomes and c-Maf. Cluster F and G likely represent exhausted states as well, since both expressed high PD-1 levels and Eomes. Cluster F didn’t express as many inhibitory receptors as cluster H, and cluster G was the only cluster that expressed KLRG1 and T-bet among clusters E-H. Recent studies suggest CD8 T cells with an effector-like profile can represent a transition state before terminal exhaustion (Beltra et al., 2020). Similarly, KLRG1 is expressed on partially exhausted cells (Bengsch et al., 2010; Long et al., 2016). In summary, we extend the results of our RNA-seq analysis by showing that helped WT CD8 T cells included subsets of cells that were effector cells with cytotoxic features. In contrast, helpless MHCII KO CD8 T cells appeared to be in various states of exhaustion.

### Validation of the “helped” and “helpless” CD8 T cell states using an independent model

We considered whether a developmental or intrinsic defect in MHCII KO CD8 T cells could explain the differences we observed between “helped” and “helpless” CD8 T cells. We analyzed splenic CD8 T cells from uninfected WT and MHCII KO mice and found higher expression of Lag-3 and PD-1 on MHCII KO CD8 T cells. (Figure S3A-C). Although these coinhibitory receptors were expressed at very low frequencies of CD8 T cells from uninfected mice compared to cells from the lungs of infected mice, it indicated that MHCII KO CD8 T cells differ from WT CD8 T cells (Do et al., 2012). Therefore, we developed an independent model to validate our findings. Purified splenic CD8 T cells from naïve C57BL/6J mice were transferred into TCRα KO recipients with or without CD4 T cells. As CD4 T cells can transfer protection, which would be a potential confounder, we sought to minimize their contribution to protection by using CD4 T cells from the P25 TCRtg mouse. P25 TCRtg CD4 T cells (hereafter, P25 cells) are specific for Ag85b, a protein that is downregulated by Mtb within weeks of infection (Bold et al., 2011; Moguche et al., 2017). The ability of P25 cells to transfer protection was largely independent of their number (Figure S4). By restricting the diversity and number of transferred CD4 T cells, and selecting an antigen that is transiently expressed, we hoped to limit the impact of CD4 T cells on protection while still providing CD4 T cell help. TCRα KO mice received 5 million CD8 T cells and 100,000 P25 cells (hereafter, helped mice) or only 5 million CD8 T cells (hereafter, helpless mice).

We compared the survival of helped and helpless mice to recipient TCRα KO mice that received only P25 cells or no cells after Mtb infection (Figure 4A). TCRα KO mice have a median survival time (MST) of 39 days. Helpless mice (CD8 T cells only) have a MST of 46 days, which shows that CD8 T cells alone conferred little protection. TCRα KO mice that only received P25 cells survived 88 days (MST). In contrast, helped mice survived 136 days (MST). We determined whether P25 and CD8 T cells synergized to promote survival using the statistical models developed by Demidenko and Miller and based on the Bliss definition of drug independence (Demidenko and Miller, 2019). The additive effect of P25 and CD8 T cells on survival (i.e., independent effect) can be calculated as 1 – (1-S_A_(t))x(1-S_B_(t)), where S_A_(t) and S_B_(t) are the survival curves of mice received P25 and CD8 T cells, respectively. The curve generated by this function largely overlaps with the survival curve of mice received P25 cells, indicating that CD8 T cells by themselves, contribute little to the overall survival (Figure 4B). The actual survival curve of helped mice is significantly greater than their predicted independent effects, establishing synergy between P25 and CD8 T cells (Figure 4B). We infer that the prolonged survival of mice that received both polyclonal CD8 T cells and P25 cells arose from the action of helped CD8 T cells.

**Figure 4.**
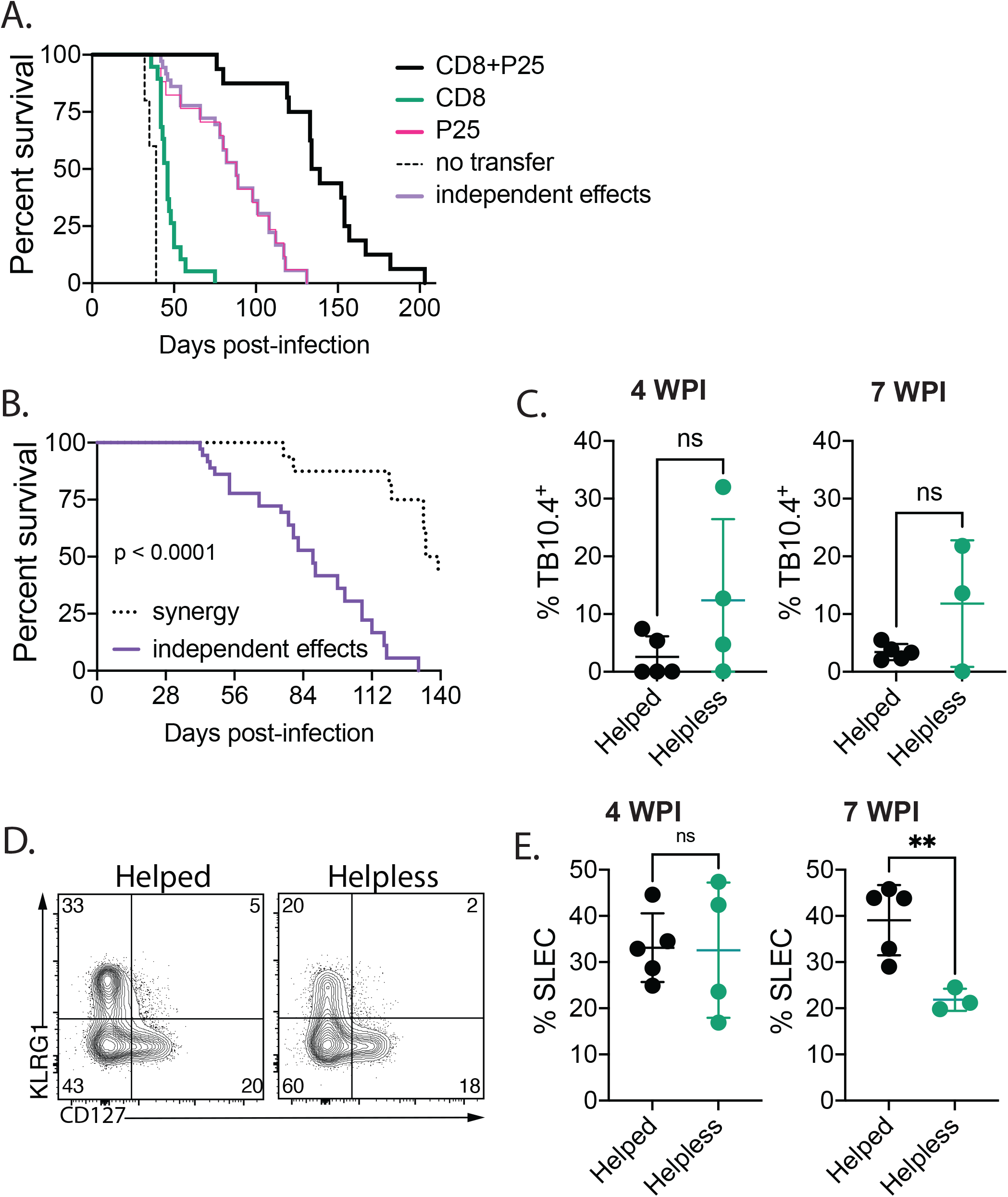
An adoptive transfer model confirms the helped vs. helpless CD8 T cell phenotype. Purified polyclonal CD8 T cells from C57BL/6J mice were transferred to TCR*α* KO mice with or without Ag85-specific CD4 T cells (P25) (i.e., helped and helpless, respectively), and then infected with aerosolized Mtb. (A) The survival of helped, helpless, TCR*α* KO mice that received only P25 cells, and TCR*α* KO mice that didn’t receive any cells was monitored. Data were combined from 3 experiments using a total of 17-19 transferred mice per group and 5 TCR*α*KO mice that received no cells. The difference between the groups was statistically significant (p<0.0001) as determined using the log-rank test for trend. (B) Determination of synergy between P25 and CD8 T cells as described in the text. The theoretical additive benefit is calculated by the “independent effects” function, while the “synergy” group is the actual survival observed after transfer of P25 and CD8 T cells (i.e., from (A)). These two scenarios differed statistically (p<0.0001) by the log-rank test. (C) Lungs from TCR*α* KO mice that received P25 and CD8 T cells (helped) vs. only CD8 T cells (helpless) were analyzed at 4 and 7 wpi by flow cytometry to determine the frequencies of TB10.4_4-11_ -specific CD8 T cells. (D) The proportion of lung KLRG1^+^CD127^−^(SLEC) and KLRG1^−^CD127^+^(MPEC) CD8 T cells was determined 7 wpi. Quadrant numbers represent percentages. (E) The frequencies of lung KLRG1^+^CD127^−^ (SLEC) among total CD8 T cells was determined 4 and 7 wpi by flow cytometry and analyzed statistically. (C, E) Bars, mean ± SD. Data are representative of three independent experiments, 3-5 mice/group. Statistical significance was analyzed by unpaired t test. p-values: **, p<0.01. ns, no significant differences.

Like the MHCII KO mice, antigen-specific CD8 T cell responses (TB10.4^+^) were generated in both helped and helpless mice (Figure 4C). Very few 32A_309-318_ -specific CD8 T cells were detected. The frequency of TB10.4_4-11_ -specific CD8 T cells was comparable between helped and helpless mice. More CD8 T cells differentiated into SLECs in helped than in helpless mice at 7 wpi (Figure 4D and 4E). In contrast, the MPEC frequencies were similar. Thus, this transfer model shows that CD8 responses to Mtb could be generated in the absence of CD4 T cells, but CD4 T cells helped CD8 T cells differentiated into SLECs and together, had a synergistic effect on survival.

### Common features of “helped” and “helpless” CD8 T cell in the WT/KO and the adoptive transfer model

We analyzed helped and helpless lung parenchymal CD8 T cells from the adoptive transfer model to determine whether they shared phenotypic characteristics with CD8 T cells from WT and MHCII KO mice. Importantly, we found that in the adoptive transfer model, helped CD8 T cells expressed more of the NK cell receptors 2B4 and NKG2D and transcription factor T-bet. In contrast, helpless CD8 T cells had higher expression of Eomes (Figure 5A).

**Figure 5.**
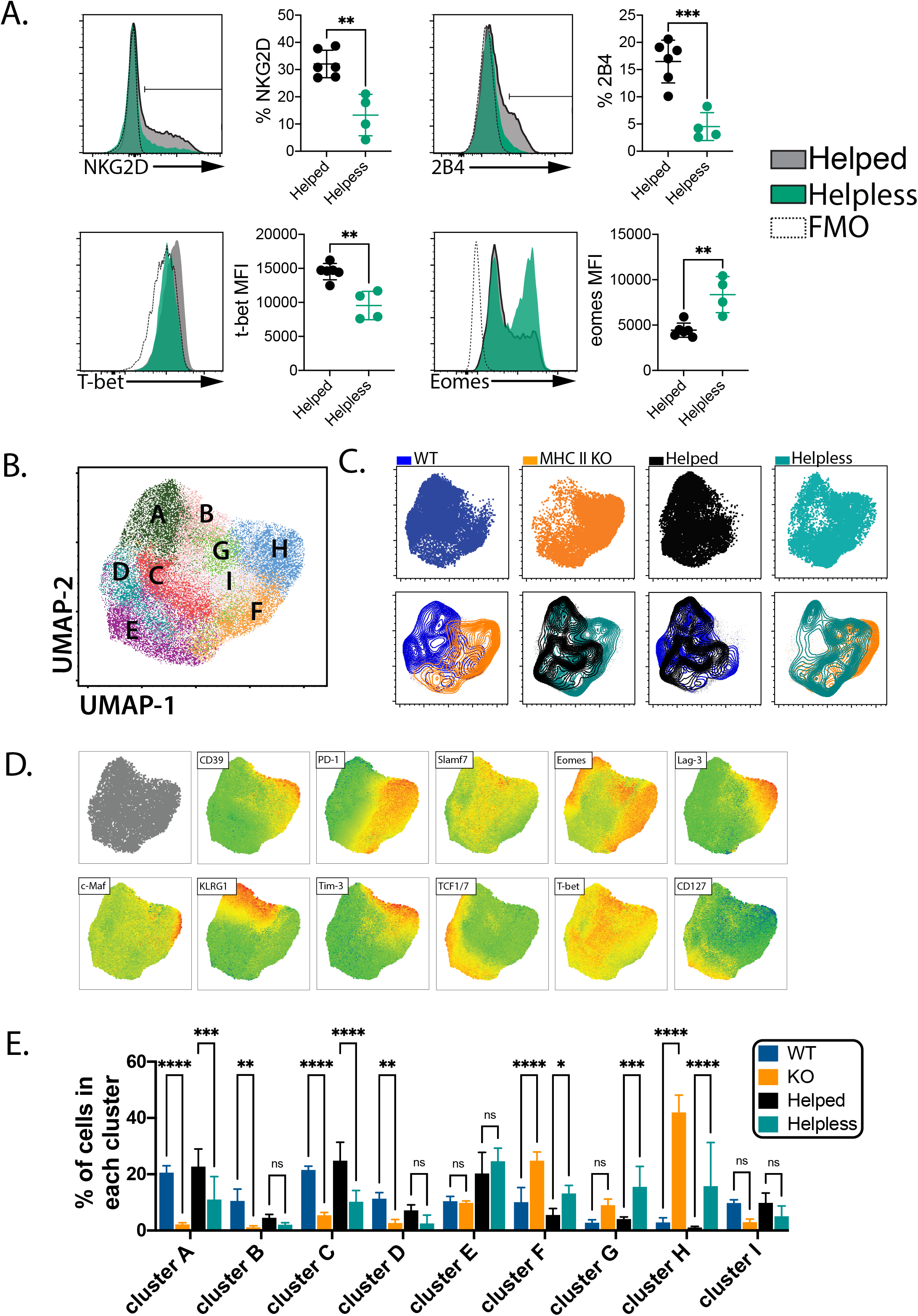
Comparison of WT/KO and adoptive transfer model finds shared “helped” and “helpless” characters. (A) Helped and helpless mice (i.e., the adoptive transfer model) were infected with aerosolized Mtb for 6 weeks. Anti-CD45-AF647 was injected intravenously 3 minutes before euthanasia. Lung parenchymal (CD45iv^−^CD44^+^CD62L^−^) CD8 T cells were analyzed by flow cytometry to determine the expression of NK receptors (percentage) and transcription factors (MFI). The histograms show representative data and the accompanying graphs represent the statistical analysis. Bars, mean ± SD. (B-E) Lung parenchymal CD8 T cells from WT/KO mice 8wpi and helped/helpless mice 6wpi were used to create UMAP projections and analyzed with PhenoGraph. (B) Nine clusters were identified with PhenoGraph analysis. (C) Pairwise comparisons of WT, KO, helped, and helpless lung parenchymal CD8 T cells were overlayed onto the UMAP projection. (D) UMAP projections were overlayed with expression of each indicated marker. (E) The percentage of each cluster among WT, KO, helped or helpless lung parenchymal CD8 T cells were compared statistically. Bars, mean ± SD. (A-E) Representative data of two independent experiments, 4-5 mice/group. Statistical significance was analyzed by unpaired t test (A) or two-way ANOVA (E). p values: **, p<0.01; ***, p<0.001; ****, p<0.0001. ns, no significant differences.

Next, we created UMAP projections and performed PhenoGraph™ analysis using WT, KO, helped, and helpless CD8 T cells (Figure 5B-E). In this analysis, we compared differences between helped vs. helpless CD8 T cells, as well as differences between the two models (i.e., WT/KO vs. adoptive transfer). The clusters were similar to those identified in the WT/KO model (Figure 3C-E), with one additional cluster (“I”, Figure 5B). Helped and helpless CD8 T cells differed in many clusters (Figure 5C-E). Helped CD8 T cells were significantly enriched in cluster A and C, which contained most of the effector cells. In contrast, helpless CD8 T cells were more abundant in cluster F, G, and H, which contained exhausted cells in various states (Figure 5D, 5E). Thus, the differences between “helped” vs. “helpless” CD8 T cells in the adoptive transfer model were concordant with those identified in the WT/KO model, indicating that the development of T cell exhaustion arises from a lack of T cell help and not from developmental defects in MHCII KO mice.

We then compared WT to helped, and KO to helpless CD8 T cells to visualize differences between these two models (Figure 5C, 5E). WT and helped CD8 T cells were similar except for an increased frequency of helped CD8 T cells in cluster E, which are progenitor exhausted CD8 T cells. Helpless CD8 T cells were also enriched in cluster E, indicating cluster E was characteristic of the adoptive transfer model. When compared to WT and helped CD8 T cells, KO and helpless CD8 T cells were significantly enriched in cluster F and H, which appear to be exhausted and terminally exhausted cells. The differences in the exhausted states of CD8 T cells from the WT/KO and adoptive transfer models might have arisen from their analysis at different time points, 8 and 6 wpi, respectively. Nevertheless, these data demonstrate shared features of helped and helpless CD8 T cells from the two models. “Helped” CD8 T cells were more likely to express NK receptors and resemble effector CD8 T cells, while “helpless” CD8 T cells were exhausted.

### CD4 T cell help promotes IFNγ and IL-2 production by CD8 T cells that recognize Mtb-infected macrophages

To gain insight into the functional differences between “helped” vs. “helpless” CD8 T cells, we measured their ability to produce cytokines when stimulated. We showed previously that TB10.4_4-11_-specific CD8 T cells did not efficiently recognize infected macrophages. As we are unaware of any class I MHC Mtb epitopes that are presented on K^b^ or D^b^ by Mtb-infected cells, we chose to use Mtb-infected macrophages as a physiological stimulus. CD8 T cells from Mtb-infected WT or MHCII KO mice (Figure 6A-C) or from Mtb-infected helped and helpless mice (i.e., the transfer model; Figure 6D-F) were purified and cultured with uninfected or Mtb-infected macrophages. The CD8 T cells from WT mice produced more IFNγ, TNF, and IL-2 than CD8 T cells from MHCII KO mice (Figure 6A-C). Similarly, CD8 T cells cotransferred with P25 cells produced more IFNγ and IL-2 than CD8 T cells transferred alone (Figure 6D-E). TNF production was comparable in the adoptive transfer model (Figure 6F). Thus, we showed that CD4 T cells provide help for CD8 T cell production of IFNγ in both of our helped/helpless CD8 T cell models using Mtb-infected macrophages as a physiological stimulus. This is consistent with previous studies that demonstrated a reduction of IFN*γ*-producing CD8 T cells in the absent of CD4 T cells (Bold and Ernst, 2012; Green et al., 2013). Additionally, we show that CD4 T cell help is critical for CD8 T cell production of TNF and IL-2. These functional changes parallel the effector vs. exhausted features that distinguish “helped” vs. “helpless” CD8 T cells in both models.

**Figure 6.**
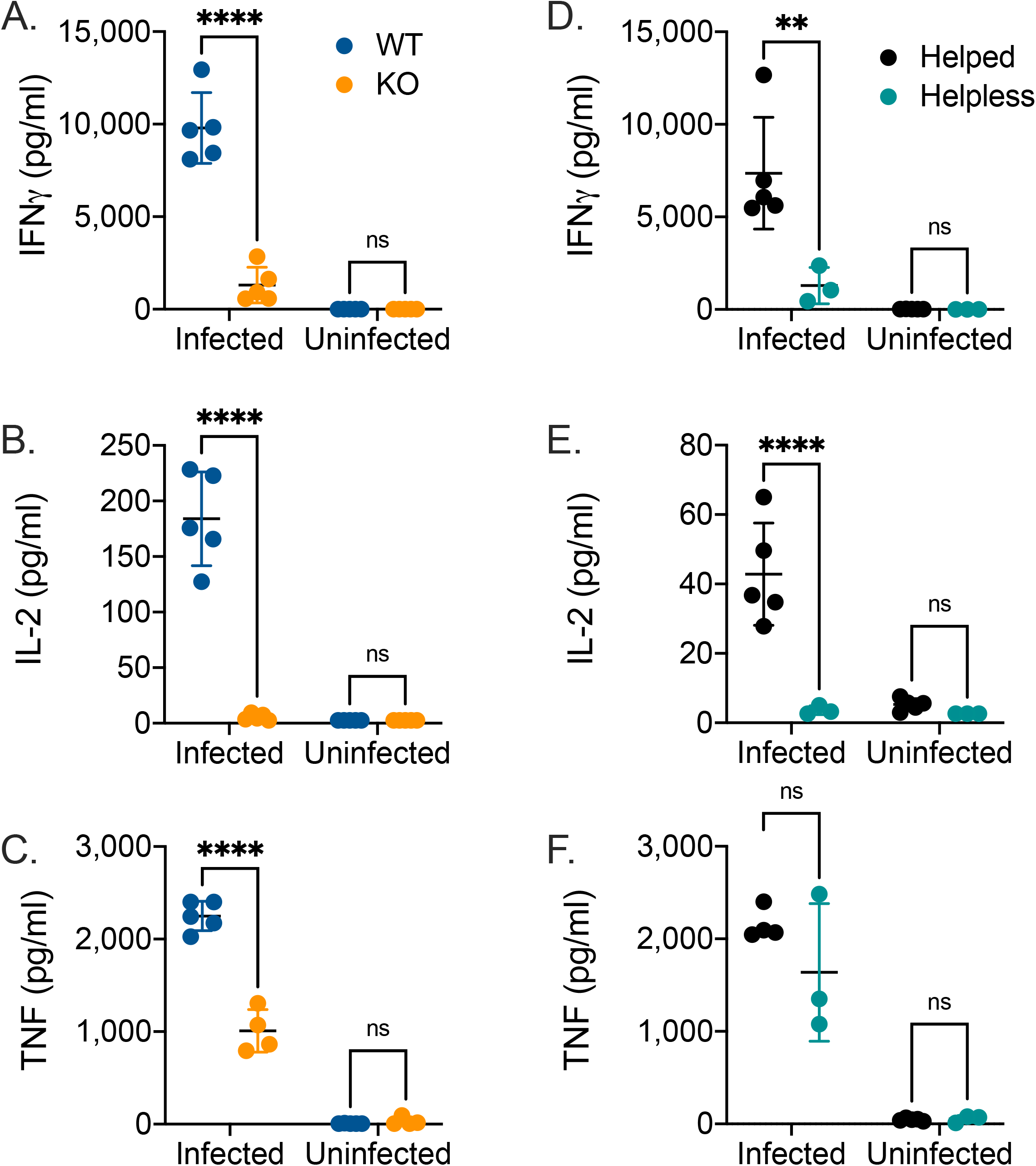
CD4 T cell help enhances CD8 T cell cytokine productions upon recognition of infected macrophages. Purified CD8 T cells from WT or MHCII KO mice 8wpi (A, B, C) or from helped or helpless mice (i.e., the adoptive transfer model) 7wpi (D, E, F) were cultured with Mtb-infected macrophages (M*ϕ*) at a CD8:M*ϕ* ratio = 1:1. IFN*γ* (A, D), IL-2 (B, E), and TNF (C, F) were determined 18-24 hours later. The data are representative of three experiments with 5 mice/group (A-C) or two independent experiments with 3-5 mice/group (D-F). Bars, mean ± SD. Statistical significance was analyzed by two-way ANOVA. p-values: **, p<0.01; ****, p<0.0001. ns, no significant differences.

### “Helped” CD8 T cells more effectively restrict Mtb growth than “helpless” CD8 T cells

Whether helped CD8 T cells mediate better protection during Mtb infection is unknown. This question is difficult to address *in vivo* because it is difficult to separate the helper function of CD4 T cells from their effector function. Therefore, we quantified the ability of purified “helped” or “helpless” CD8 T cells to restrict intracellular bacterial growth *in vitro*. Using this approach, we found CD8 T cells from the lungs of WT mice inhibited Mtb growth effectively, even when the CD8: macrophage ratio was as low as 1:125 (Figure 7A), although the potency varied between experiments. Bacterial control was MHCI-restricted since CD8 T cells were able to restrict bacterial growth in infected WT macrophages but not in infected K^b^D^b^ KO macrophages (Figure 7B). WT CD8 T cells inhibited Mtb growth significantly better than MHCII KO CD8 T cells (Figure 7C). Interestingly, the colony forming unit (CFU) recovered after adding WT CD8 T cells was sometimes less than the input (i.e., day 1), suggesting that CD8 T cells are capable of promoting killing of Mtb under some conditions. These results were confirmed using CD8 T cells purified from Mtb-infected helped and helpless mice (i.e., the transfer model). Again, helped CD8 T cells inhibited bacterial growth significantly better than helpless CD8 T cells (Figure 7D). The ability of CD8 T cells to control bacterial growth depended on when they were isolated from mice. As MHCII KO mice survive longer after Mtb infection than T cell deficient mice, CD8 T cells must be able to mediate some protection *in vivo*. We hypothesized that CD4 T cell help is important in maintaining the function of CD8 T cells and preventing exhaustion. Therefore, we compared the ability of WT and MHCII KO CD8 T cells to restrict Mtb growth when isolated from mice at 5 vs. 8 wpi. MHCII KO CD8 T cells obtained after 5 wpi were able to inhibit intracellular bacterial growth while CD8 T cells isolated 8 wpi had lost their ability to restrict Mtb growth (Figure 7E). WT CD8 T cells isolated at both timepoints similarly controlled bacterial growth. In order to compare experiments where Mtb grew differently, the growth from day 1 to endpoint was normalized and the percent inhibition by CD8 T cells was calculated. The 100% inhibition indicated no Mtb growth, while 0% inhibition indicated Mtb grew similarly as in macrophages without CD8 T cells (Figure 7F). This correlated with greater inhibitory receptor expression on MHCII CD8 T cells 8 wpi (Figure 7G). These data show that “helped” CD8 T cells restricted Mtb growth more efficiently than “helpless” CD8 T cells, and this effect is due to a loss of function of “helpless” CD8 T cells late during infection.

**Figure 7.**
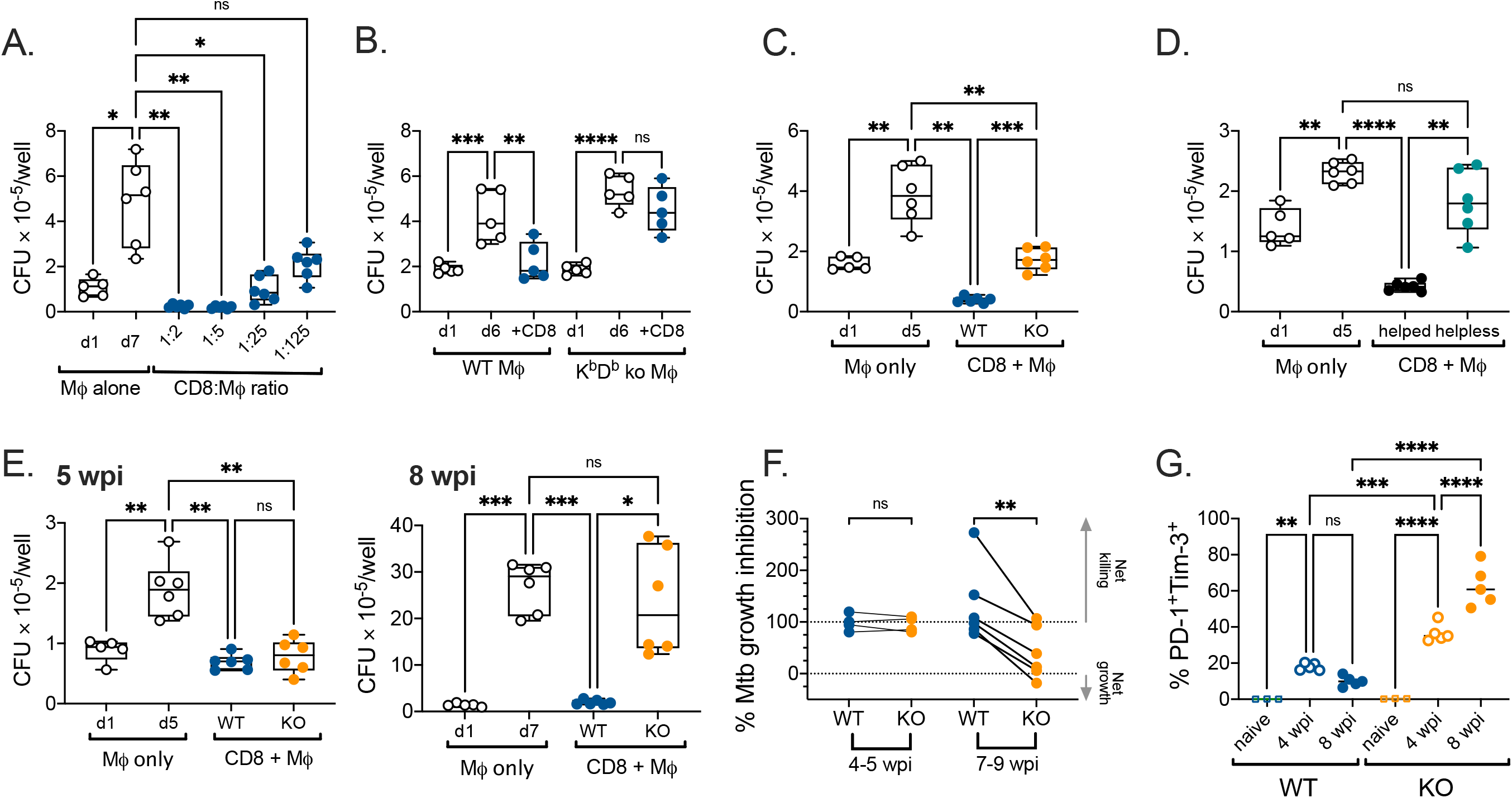
“Helped” CD8 T cells are better at Mtb control than “helpless” CD8 T cells. Purified CD8 T cells from WT or MHCII KO mice at 8 wpi (A-C, and E), or purified CD8 T cells from helped or helpless mice (i.e, the adoptive transfer model) at 7 wpi (D) were cultured with Mtb infected macrophages (M*ϕ*) at a CD8:M*ϕ* ratio = 1:2 unless otherwise indicated. Macrophages were lysed 4-6 days later and the colony forming unit (CFU) per well was determined. (A) Purified WT CD8 T cells were cultured with macrophages at different T cell to macrophage ratios. (B) Purified WT CD8 T cells were cultured with WT or K^b^D^b^ macrophages. (C) Purified WT or MHCII KO CD8 T cells were cultured with macrophages. (D) Purified helped or helpless CD8 T cells were cultured with macrophages. (E) Purified WT or MHCII KO CD8 T cells from 5 or 8 wpi were cultured with macrophages. (F) Percent inhibition by CD8 T cells from early and late timepoints was calculated. The 100% inhibition indicated no Mtb growth since day 1, while 0% inhibition indicated Mtb grew similarly to conditions that had no T cells. (G) Frequencies of PD-1^+^Tim-3^+^ CD8 T cells from spleens of naïve and lungs of 4 and 8 wpi mice. Bar, mean. (A-E) The whiskers of the Box and whisker plots indicate the maximum and minimum. Representative of 3 (A, D), 2 (B, G),6 (C), or 4 (E) experiments. Statistical significance was analyzed by one-way ANOVA. p-values: **, p<0.01; ***, p<0.001; ****, p<0.0001. ns, no significant differences.

## Discussion

The relative importance of CD4 and CD8 T cells for immunity to Mtb continues to be debated. Tackling this question is confounded by several issues. CD4 and CD8 T cells differ in the way they are activated. CD8 T cells survey the cytosol, which is sampled by MHCI, while CD4 T cells survey antigens produced in endosomal vesicles or acquired by endocytosis and sampled by MHCII. Thus, assaying T cell responses by in vitro stimulation with exogenous antigen (e.g., PPD or Mtb lysate) biases the results towards CD4 T cells. Optimally stimulating CD8 T cells requires defined epitopes or delivering the antigen into the cytosol. Thus, many immunogenicity studies find vaccines elicit primarily CD4 T cells responses, but this could be prejudiced by the stimulation conditions. Consequently, many investigators focus on vaccine strategies that elicit CD4 T cell effector responses. We hypothesize the potential of CD8 T cells to mediate protection is underestimated because they require CD4 T cell help to optimally express effector functions that inhibit Mtb growth.

We used RNA-seq to detect differences between WT and MHCII KO CD8 T cells during Mtb infection in the mouse model. These “helped” vs. “helpless” features were confirmed by flow cytometry in two independent models. Among the helped CD8 T cells, there were higher frequencies of SLECs and MPECs, and cells that expressed NK receptors. This may explain why CD4 T cells enhanced CD8 T cell lysis of Mtb-infected macrophages (Serbina et al., 2001). In contrast, helpless CD8 T cells resembled exhausted cells. We measured cytokine productions by CD8 T cells in response to Mtb-infected macrophages. Helped CD8 T cells secreted more IFNγ, TNF, and IL-2, while helpless CD8 T cells produced less cytokines. Previous studies found CD4 T cells enhance Mtb-specific CD8 T cell production of IFNγ (Bold and Ernst, 2012; Green et al., 2013), and our data suggest that maintenance of cytokine production depends on CD4 T cells preventing CD8 T cell exhaustion.

Since CD4 T cells mediate protection during Mtb infection, it’s difficult to study CD4 T cell help to CD8 T cells without the confounder of direct protection by CD4 T cells. Because of this, none of the previous studies addressed whether CD4 T cell help is necessary for protection mediated by CD8 T cells. We developed a novel adoptive transfer model to identify characteristics of helped and helpless CD8 T cell responses by using P25 TCRtg CD4 T cells that provide help but only limited protection. The combination of polyclonal CD8 T cells plus P25 CD4 T cells led to a synergistic effect in the survival of Mtb-infected mice. Finally, lung CD8 T cells purified from both of our models and cultured with infected macrophages showed that helped CD8 T cells directly restricted Mtb growth more efficiently than helpless CD8 T cells. Without CD4 T cells in vivo, mice succumb prematurely to TB despite lung IFNγ levels remaining elevated (Caruso et al., 1999). Thus, it is uncertain whether greater CD8 T cells production of IFNγ in vivo would correlate with protection in the absence of CD4 T cells. These results highlight the requirement of CD4 T cells for protective CD8 T cell responses and indicate that compromised CD8 immunity likely contributes to the early death of mice lacking CD4 T cells.

An underappreciated role of CD4 T cells is to help CD8 T cells resist exhaustion, a state characterized by the expression of multiple co-inhibitory receptors and accompanied by the gradual loss of effector functions. Exhausted CD8 T cells express a unique transcriptional network that inhibits their differentiation into effector or memory cells (Angelosanto et al., 2012; Chen et al., 2019b). How CD8 T cell exhaustion contributes to TB pathogenesis is not completely clear; nevertheless, T cells with features of exhaustion are detected during TB both in mice and in people (Jayaraman et al., 2016). Why helpless CD8 T cells become exhausted during the early stages of infection is unknown. One hypothesis is antigen persistence during chronic infection leads to persistent TCR engagement and exhaustion, supported by the finding that the transcription factors downstream of TCR signaling, including IRF4, NFAT, Nr4a1, Nr4a2, and Nr4a3, enhance exhaustion (Chen et al., 2019a; Man et al., 2017; Martinez et al., 2015). Alternatively, suboptimal priming of CD8 T cells in the absence of CD4 help favor the formation of exhausted progenitors rather than effector cells (Busselaar et al., 2020; Kanev et al., 2019; Snell et al., 2018). Our results are better explained by the chronic antigen stimulation model. We found a difference between WT and MHCII KO mice in inhibitory receptor expression and lung CFU by 4 wpi. We speculate that the higher bacterial burden in MHCII KO mice leads to CD8 T cell exhaustion.

An area for further investigation is the molecular nature of help signals during TB. When CD4 T cells recognize antigen presenting cells (APC), CD40/CD40L interaction, combined with innate signaling by pattern recognition receptors, leads to DC activation, cytokine production and upregulation of co-stimulatory molecules (Bennett et al., 1998; Schoenberger et al., 1998). Cytokines including IL-12, IL-15, and type I interferon, and co-stimulatory ligands CD70 and CD80/86, are the molecular basis of “help” that DCs relay to CD8 T cells (Laidlaw et al., 2016). In addition, CD4 T cells produce IL-2 and IL-21, which promote CD8 T cell survival and acquisition of effector function (Wilson and Livingstone, 2008). Thus, CD4 T cell help CD8 T cells both directly and indirectly. IL-2 treatment also restored T cell responses in a mouse model, in which Mtb proteins were repeatedly administered to study T cell dysfunction (Liu et al., 2019). CD8 T cells require IL-21 to control chronic viral infection (Elsaesser et al., 2009; Fröhlich et al., 2009; Yi et al., 2009), and we showed previously that IL-21 receptor deficiency impairs CD8 T cell responses during TB and leads to increased Tim-3 and PD-1 expression (Booty et al., 2016). Whether exhaustion of helpless CD8 T cells develops because of reduced IL-2 or IL-21 levels needs further investigation. An interesting clue is that at 5 wpi, both helped and helpless CD8 T cells restricted Mtb growth in vitro; however, the function of helpless CD8 T cells obtained later during infection seemed to decline. This may indicate that “help” is needed to maintain effector function or prevent T cell exhaustion.

Mtb-specific CD8 T cell responses are detected in infected people and animal models and are recognized as an integral part of immune response to Mtb. CD8 T cells were considered to play a minor role compared to CD4 T cells based on only moderate reduction in the survival of mice with defective CD8 T cell responses compared to normal mice. Here, using a variety of approaches including two independent in vivo models, we unequivocally show that CD4 T cells play an essential role in helping CD8 T cells develop into effector T cells that can restrict Mtb growth. These data are consistent with a greater appreciation for the role of CD8 T cells in primary Mtb infection and Mtb-challenge after vaccination in the non-human primate model (Chen et al., 2009; Hansen et al., 2018). We suggest that vaccine strategies that enhance synergy between CD4 and CD8 T cells would be more effective at eliciting protective immunity.

## Acknowledgements

We thank Dr. Eugene Demidenko (Dartmouth University, Hanover, NH) for his help on the drug-independence model, and I-Hao Wang for help with the RNA-seq analysis using R. This work was funded by NIH/NIAID grants to S.M.B.: R01AI106725 and P01 AI073748.

## Author contribution

Conceptualization, Y.L. and S.M.B.; Methodology, Y.L., P.B.S., and S.M.B.; Investigation, Y.L., P.B.S., S.B., J.P., and K.C.; Formal analysis, Y.L. and S.M.B.; Writing-Original Draft, Y.L.; Writing-Review & Editing, Y.L., P.B.S., S.B., J.P., K.C., and S.M.B.; Supervision, S.M.B.; Funding Acquisition, S.M.B.

## Declaration of interests

The authors declare no competing interests.

## STAR Methods

### Resources availability

#### Lead contact

Further information and requests for resources and reagents should be directed to and will be fulfilled by the Lead Contact, Samuel Behar.

#### Material availability

This study did not generate unique reagents.

#### Data and code availability

All RNASeq data are available from the GEO database (accession number: pending).

### Experimental Model and Subject Details

#### Mice

P25 TCRtg, MHCII KO, and TCR*α* KO mice were purchased from Jackson Lab and bred locally. C57BL/6J mice were purchased from Jackson Lab. K^b^D^b^ KO mice were a generous gift from Dr. Kenneth Rock (University of Massachusetts Medical School) (Pérarnau et al., 1999). Experimental mice were 7-12 weeks old and sex-matched. In adoptive transfer experiments, 7-12 weeks old, sex-matched TCR*α* KO mice from different litters were randomly assigned for receiving P25, CD8, or both cells. All animal studies were conducted using the relevant guidelines and regulations, and approved by the Institutional Animal Care and Use Committee at the University of Massachusetts Medical School (UMMS) (Animal Welfare A3306-01), using the recommendations from the Guide for the Care and Use of Laboratory Animals of the National Institutes of Health and the Office of Laboratory Animal Welfare.

#### Mtb strain

The Erdman strain of *Mycobacterium* tuberculosis, which has been passaged through mice, was used for aerosol infection as described (Chackerian et al., 2001). H37Rv or CDC1551 strain was used for *in vitro* infection of macrophages. CDC1551 was provided by Dr. Christopher Sassetti (University of Massachusetts Medical School)

### Method Details

#### Aerosolized Mtb infection of mice

Mice were infected by the aerosol route as previously described (Lee et al., 2020). Frozen bacterial stocks were thawed and sonicated for 1 minute and then diluted into 5 ml of 0.01% Tween-80 in PBS. The diluted bacterial suspension was aerosolized to infect mice using a Glas-Col chamber (Terre Haute). The average number of bacteria delivered into the lung was determined for each experiment by plating lung homogenate from 4-5 mice 24 hours after infection and ranged between 40-150 CFU/mouse.

#### Survival studies

Infected mice were monitored weekly in accordance with IACUC guidelines using the Body Condition Score (BCS) and serial weight determinations. Mice with a BSC score less than or equal to 2 or had lost 20% body weight were euthanized.

#### Bacterial burden in lung and spleen

Infected mice were euthanized at pre-determined timepoints, and the left lung or whole spleen were homogenized using 2 mm zirconium oxide beads (Next Advance) in a FastPrep homogenizer (MP Biomedicals). Tissue homogenates were serially diluted and plated on 7H11 agar plates (Hardy Diagnosis). CFU was enumerated after 19-21 days of incubation at 37°C and 5% CO_2_.

#### Lung cell preparation

Lungs were perfused with 10 ml of cold RPMI (Gibco) before removal. Single cell suspensions were prepared by homogenizing lungs using a GentleMACS tissue dissociator (Miltenyi), digesting with 300 U/ml collagenase (Sigma) in complete RPMI (10% FBS, 2 mM L-Glutamine, 100 units/ml Penicillin/Streptomycin, 1 mM Na-Pyruvate, 1X Non-essential amino acids, 0.5X Minimal essential amino acids, 25 mM of HEPES, and 7.5 mM of NaOH) at 37°C for 30 minutes, and followed by a second run of dissociation using the GentleMACS. Suspensions were then sequentially filtered through 70 µm and 40 µm strainers.

#### Intravascular staining

Mice were injected intravenously with 0.2 ug/mouse of anti-CD45-AF647 in 200 ul of injection medium (2% FBS in PBS) 2 minutes before euthanizing with CO_2_. Lungs were then perfused and removed about 3 minutes after anti-CD45-AF647 injection. Lymphocytes from blood collected in RPMI containing 40 U/ml of heparin were isolated using Lympholyte*®*(CEDARLANE), and analyzed by flow cytometry to confirm uniform staining with anti-CD45-AF647.

#### Flow cytometric analysis

Cells were first stained with Zombie Fixable Viability dye (Biolegend) for 10 minutes at room temperature (RT). For surface staining, cells were stained with antibodies in autoMACS running buffer (Miltenyi) containing 5 ug/ml of anti-mouse CD16/32 (BioXcell) for 20 minutes at 4°C. In some experiments, TB10.4_4-11_ and/or 32A_309-318_ tetramers were used with other antibodies. To measure transcription factor (TF) expression, cells were fixed and permeabilized for 30 minutes at RT, followed by staining for 30 minutes with antibodies to the different TF at RT using the Foxp3/TF staining buffer set (ThermoFisher). Samples were acquired on Aurora (Cytek) or MACSquant (Miltenyi). Flow data were analyzed using FlowJo v10.7.1, and FlowJo plugins UMAP v3.1 and PhenoGraph v3.0. For UMAP projections of WT/KO CD8 T cells (i.e., Fig 3C and S2C), 2000 CD45iv^−^CD44^+^CD62L^−^ CD8 T cells/mouse from 5 WT and 5 MHCII KO mice were concatenated to one FCS file before creating UMAP projections. In UMAP projections comparing two models (i.e., Fig 5D), 2000 CD45iv^−^CD44^+^CD62L^−^ CD8 T cells/mouse from 5 WT and 5 KO, and 6 helped and 4 helpless mice from adoptive transfer model were concatenated and created UMAP projections. K= 300 and K= 400 were used in PhenoGraph analysis in Fig 3D and Fig 5B, respectively, and the resulting clusters were overlayed onto UMAP projections.

#### Adoptive transfer model

Spleens and lymph nodes from C57BL/6J mice were mechanically disrupted onto 70 µm strainers using the plungers of 3 ml syringes. CD8 T cells were then purified from cell suspensions using CD8a (Ly-2) microbeads (Miltenyi) and autoMACS separator (Miltenyi). Preparing cell suspension with same method, CD4 T cells were purified from spleens and lymph nodes of P25 mice using MojoSort^TM^ CD4 isolation kit and magnet (Biolegend). Purities of cells were determined for each experiment using flow cytometry. 2-5 million of CD8 T cells were transferred into TCR*α* KO mice with or without distinct congenic marked 10^5^ P25 cells before infecting with Erdman.

#### *In vitro* CFU assay

Purified CD8 T cells were cultured with infected macrophages to quantify their ability to inhibit Mtb growth. The assay is described in following sections:

##### Preparation of Mtb

CDC1551 was grown in 7H9 media (supplemented with 10% OADC, 0.05% of Tween-80, and 0.2% glycerol) until an OD_600_ = 0.6-1, washed with RPMI, and opsonized with TB coat (RPMI 1640 containing 1% heat-inactivated FBS, 2% human serum, and 0.05% Tween-80). The bacteria were passed through 5 µm filter to remove clumps and then enumerated by microscopy using a Petroff-Hausser counting chamber. The concentration was adjusted to provide a multiplicity of infection (MOI) of 2.

##### Macrophage collection and infection

Thioglycolate-elicited peritoneal macrophages (TG-PMs) were obtained by peritoneal lavage 3-5 days after intraperitoneal injection of mice with 3% thioglycolate, and purifying from the lavage using CD11b microbeads (Miltenyi) and LS columns (Miltenyi). Purified TG-PMs were plated in Nunc^TM^ Up-Cell^TM^ 12-well plates (10^6^/well), and once adhered, infected with CDC1551 overnight. Infected TG-PMs were detached and washed with cold complete RMPI, and re-seeded as 10^5^/well in 96-well flat bottom plate before CD8 T cells were added.

##### CD8 T cell isolation and co-culture

CD8 T cells were purified from the lungs of infected mice at indicated timepoints using CD8a (Ly-2) microbeads (Miltenyi) and an autoMACS separator (Miltenyi). The purity of cells was determined for each experiment and was typically 95%. Purified CD8 T cells were added to infected TG-PMs at a ratio 1: 2 (CD8 T: TG-PM) or as indicated. TG-PMs were lysed with 1% Triton X-100 after 4-6 days of co-culture, and CFU was determined by plating serial dilutions of the lysate on 7H10 or 7H11 plates (Hardy Diagnosis). Percent inhibition by CD8 T cells was calculated as: 100 x (Mtb net growth without CD8 T cell – Mtb net growth with CD8 T cell)/ Mtb net growth without CD8 T cell.

#### Cytokine production by CD8 T cells

Lung CD8 T cells were isolated and cultured with macrophages infected with CDC1551 or H37Rv, as described above with the modifications that an MOI=5 was used and CD8 T cells were added into infected TG-PMs at a 1:1 ratio. Culture supernatants were collected after 18-24 hours of co-culture, and IL-2, IFN*γ*, and TNF were assayed using LEGENDplex^TM^ (Biolegend).

#### RNA-seq

CD8 T cells were collected and purified from lungs of infected mice 8wpi as described above. Purified CD8 T cells from individual lungs were suspended in RNAprotect® cell reagent (Qiagen) and the RNA extracted using RNeasy® mini kit (Qiagen). RNA samples were sequenced by GENEWIZ. Reads were aligned to the mouse genome and differential expression analysis performed by GENEWIZ. Gene Ontology analysis was performed with genes filtered with p-adjusted value (padj) < 0.05 and |log_2_ fold-change|*≥*1, or filtered with padj < 0.05 independently of fold-change using R (v 3.6.1) package clusterProfiler (v3.14.3) (Yu et al., 2012). The top 10 enriched pathways with the highest padj value were presented. GSEA was performed using the Broad institute software and published datasets of CD8 transcriptome from Gene Expression Omnibus (GEO) (Liberzon et al., 2011; Subramanian et al., 2005). Results comparing with datasets GSE30962 and GSE9650 were shown.

### Quantification and Statistical Analysis

Statistical analysis was performed using Graphpad Prism 8. P-values were calculated using unpaired t test, one-way ANOVA, or two-way ANOVA as indicated in the figure legends.

## Supplementary figure legends

**Figure S1.**
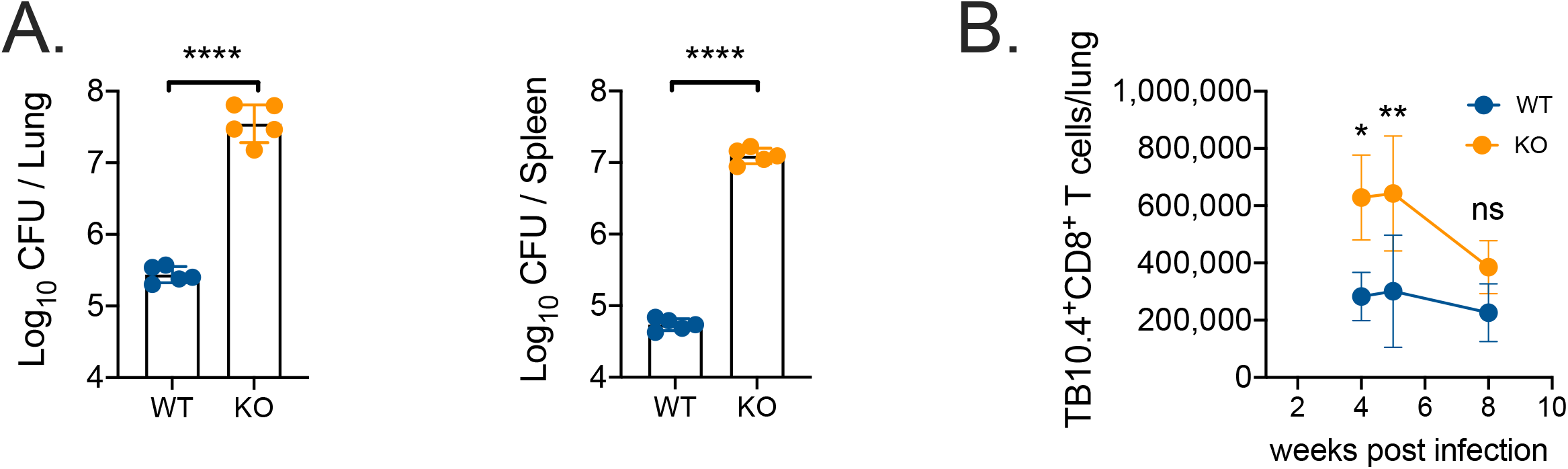
MHCII KO mice have higher bacterial burden and recruit more CD8 T cells to lungs after infection. WT or MHCII KO mice were infected with aerosolized Mtb and the bacterial burden in lungs and spleens were determined 8 wpi (A), and total TB10.4_4-11_ tetramer^+^ CD8 T cell number in lungs at 4, 6, and 8 wpi (B). Data are representative of two independent experiments, 5 mice/group. Bars, mean ± SD. Statistical significance was analyzed by unpaired t test. p-values: **, p<0.01; ****, p<0.0001.

**Figure S2.**
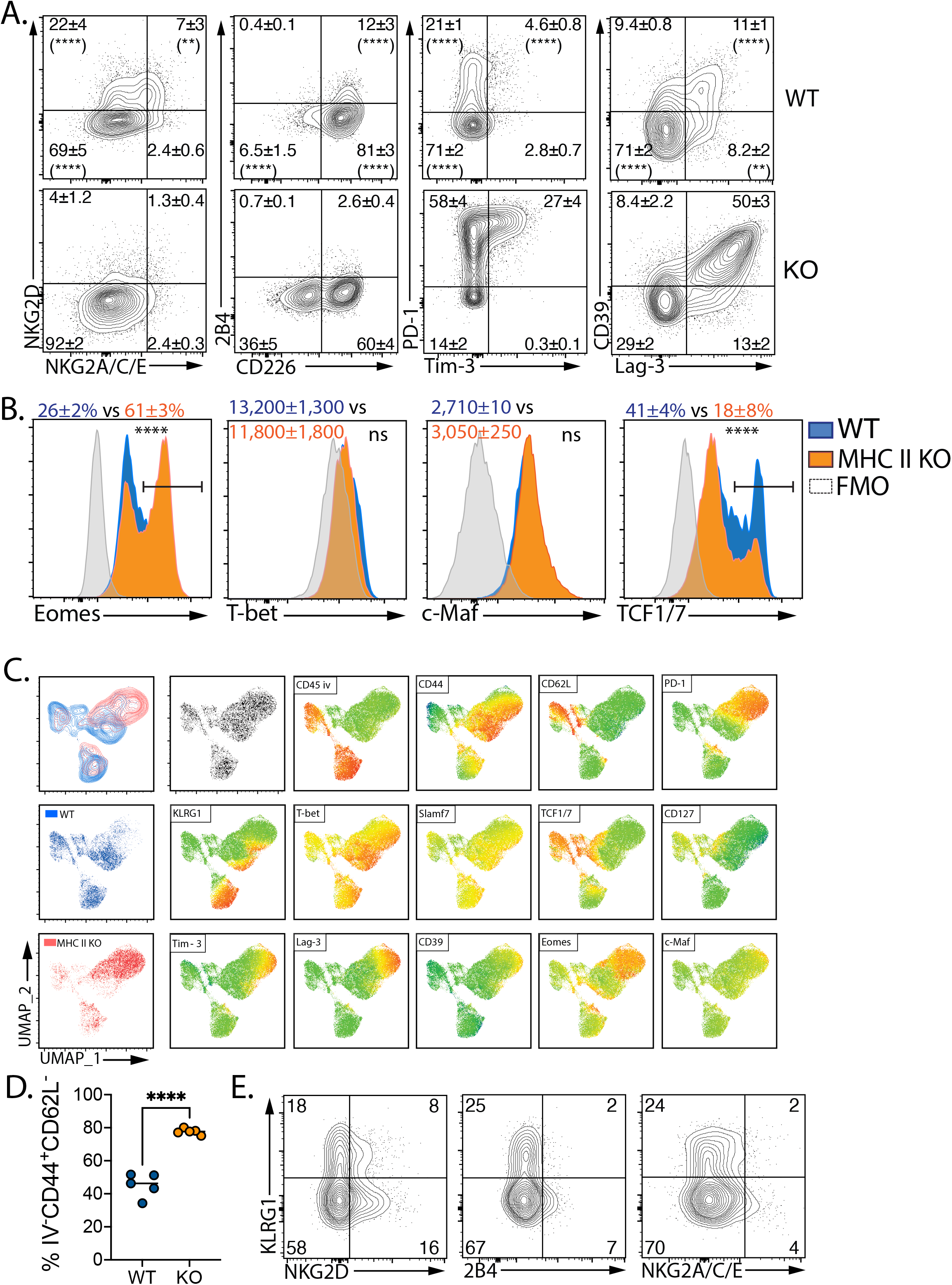
Phenotypic differences between WT and MHCII KO lung CD8 T cells. (A) WT and MHCII KO mice were infected with aerosolized Mtb and total lung CD8 T cells were analyzed by flow cytometry at 8 wpi to determine the percentage of NK and inhibitory receptors. The mean % ± SD for each quadrant is indicated (n=5/group). The percentage of WT vs. KO cells in each quadrant was compared, and if statistically significant, is indicated in the WT quadrant. (B) The median fluorescence intensity (MFI) or average percentage of cells expressing each transcription factors, ± SD, are indicated. (C) UMAP projections of lung CD8 T cells were created and overlayed with heatmaps of the indicated markers. (D) Frequencies of CD8 T cells located in parenchyma were analyzed statistically. Bar, mean. (E) Frequencies of KLRG1 and NKG2D, NKG2A/C/E, or 2B4 in lung parenchymal (CD45iv^−^CD44^+^CD62L^−^) CD8 T cells were determined. (A-E) Representative data of two independent experiments, 5 mice/group. Statistical significance was analyzed by two-way ANOVA (A) or unpaired t tests (B and D). p-values: **, p<0.01; ****, p<0.0001. ns, no significant differences.

**Figure S3.**
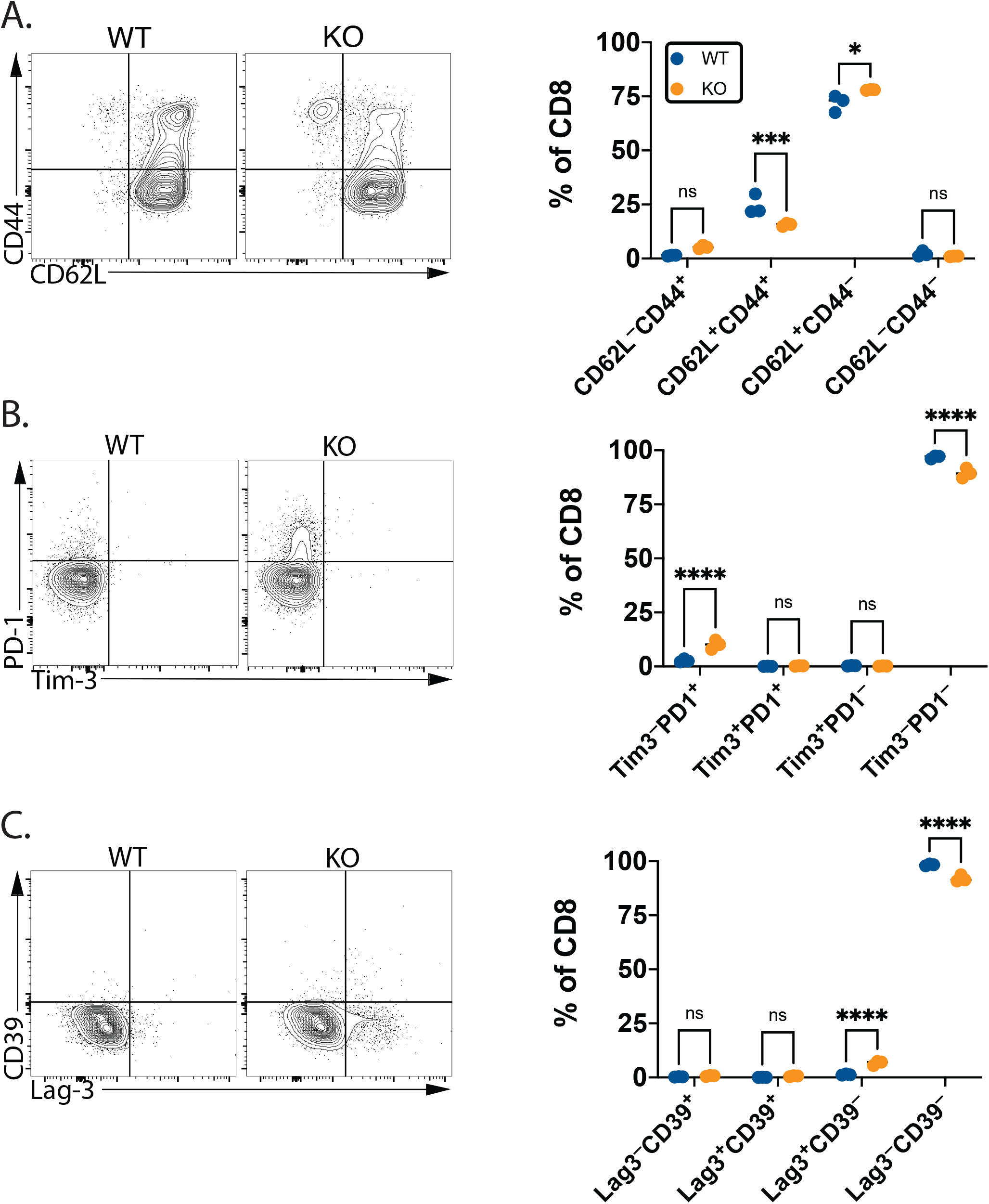
Phenotypic differences between naïve splenic WT and MHCII KO CD8 T cells. CD8 T cells in spleens of naïve WT or MHCII KO mice were analyzed by flow cytometry. (A) Expressions and percentages of CD44^+^CD62L^−^, CD44^+^CD62L^+^, CD44^−^CD62L^+^, and CD44^−^CD62L^−^ cells were determined. (B) Expressions and percentages of PD-1^+^Tim-3^−^, PD-1^+^Tim-3^+^, PD-1^−^Tim-3^+^, and PD-1^−^Tim-3^−^ cells were determined. (C) Expression and percentages of CD39^+^Lag-3^−^, CD39^+^Lag-3^+^, CD39^−^Lag-3^+^, and CD39^−^Lag-3^−^ are shown. (A-C) Bar, mean. Statistical significance was analyzed by two-way ANOVA. p-values: *, p<0.05; ***, p<0.001; ****, p<0.0001. ns, no significant differences.

**Figure S4.**
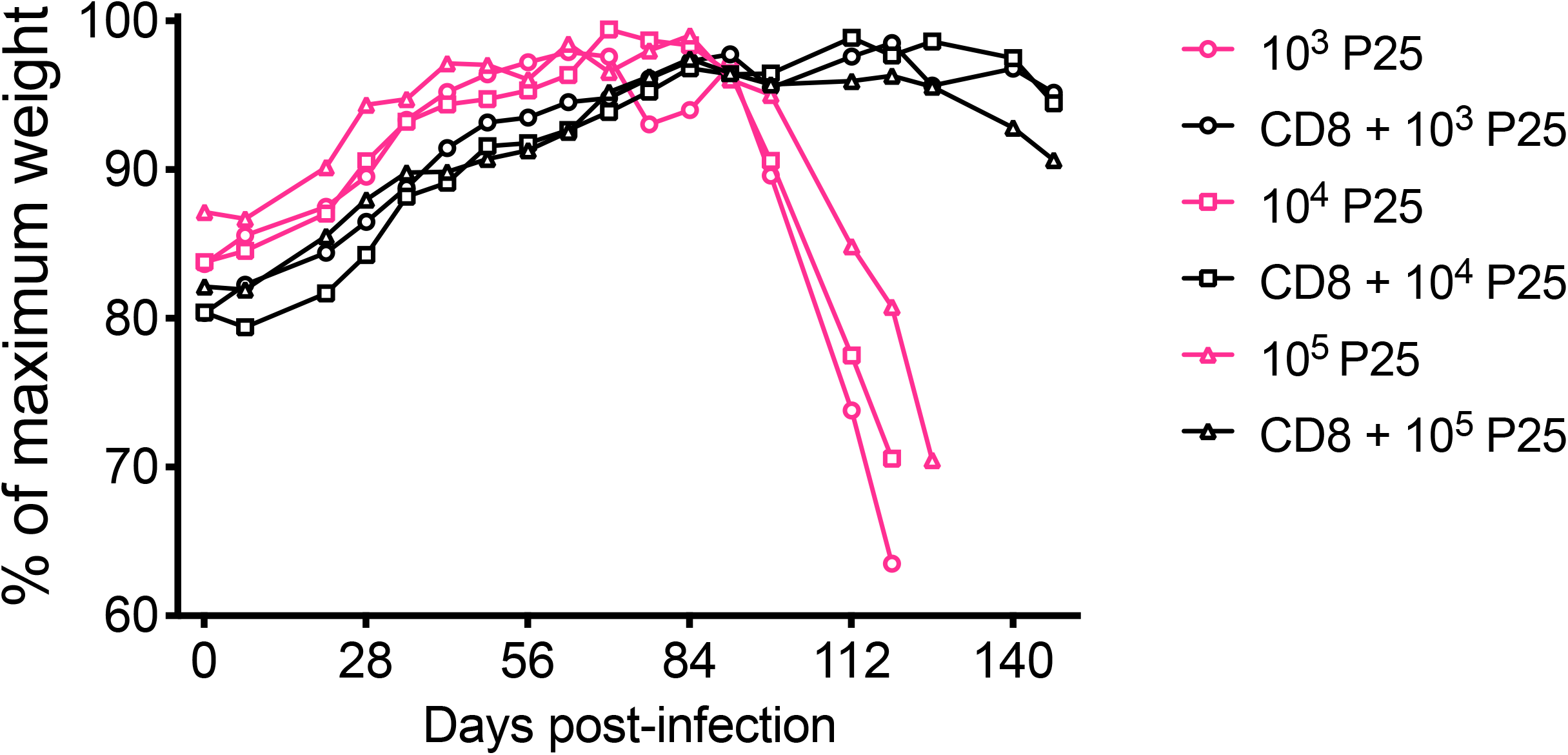
Titration of P25 CD4 T cells. Different numbers of P25 CD4 T cells were transferred, with or without 5 million polyclonal CD8 T cells, to TCRa ko mice. Mice were weighed weekly.

## Notes

### Competing Interest Statement

The authors have declared no competing interest.

